# AlbuMAX supplemented media induces the formation of transmission-competent *P. falciparum* gametocytes

**DOI:** 10.1101/2024.04.05.588064

**Authors:** Wouter Graumans, Alex van der Starre, Rianne Stoter, Geert-Jan van Gemert, Chiara Andolina, Jordache Ramjith, Taco Kooij, Teun Bousema, Nicholas Proellochs

**Affiliations:** Department of Medical Microbiology, Radboud University Nijmegen Medical Centre, Nijmegen, The Netherlands; Department of Immunology and Infection, London School of Hygiene and Tropical Medicine, London, UK

**Keywords:** Malaria, Fatty acids, Gametocyte commitment, Mosquito membrane feeding

## Abstract

Asexual blood stage culture of *Plasmodium falciparum* is routinely performed but reproducibly inducing commitment to and maturation of viable gametocytes remains difficult. Culture media can be supplemented with human serum substitutes to induce commitment but these generally only allow for long-term culture of asexual parasites and not transmission-competent gametocytes due to their different lipid composition. Recent insights demonstrated the important roles lipids play in sexual commitment; elaborating on this we exposed ring stage parasites (20-24 hours hpi) for one day to AlbuMAX supplemented media to trigger induction to gametocytogenesis. We observed a significant increase in gametocytes after AlbuMAX induction compared to serum. We also tested the transmission potential of AlbuMAX inducted gametocytes and found a significant higher oocyst intensity compared to serum. We conclude that AlbuMAX supplemented media induces commitment, allows a more stable and predictable production of transmittable gametocytes than serum alone.

**Highlights:** Gametocytes are formed when asexual parasites commit to sexual differentiation.

Sexual commitment can be promoted by environmental stressors in media formulations.

Short exposure of young asexual parasites to the serum substitute AlbuMAX achieves high proportion of committed gametocytes that are transmission-competent.

## Main text

Malaria is a mosquito-borne disease that continues to place a heavy burden on endemic countries. The parasites biomass of *Plasmodium* parasites increases by erythrocytic asexual replication and only a small subset commits to sexual stages (gametocytes) that are necessary to mediate transmission. Male and female gametocytes are formed by induction of sexual commitment, which is regulated by activation of Apatella2-g (PfAP2-G) and under epigenetic control of heterochromatin protein 1 (PfHP1) and gametocyte development 1 (GDV1) [1, 2]. Parasites commit either in the asexual cycle preceding sexual differentiation or early after merozoite invasion [3].

*Plasmodium falciparum* is the deadliest human malaria parasite, each year responsible for over half a million deaths [4], and is the only species that is regularly cultured continuously *in vitro*. Culture medium is supplemented with human serum or more easy obtainable and financially attractive serum substitutes such as AlbuMAX [5]. These substitutes can have long-term effects on parasite viability and can alter gene expression profiles, virulence factors, and drug sensitivity [6-9]. Furthermore, serum substitutes typically only allow for culture of asexual parasites and not of transmission-competent gametocytes that require different medium composition and concentration of fatty acids [10, 11]. There are several approaches to induce commitment to gametocytogenesis (reviewed in [10]) that are largely based on the principle that commitment is triggered by parasite stress, with recent developments revealing the critical role on fatty acids in sexual conversion [11-15]. The aim of the current study was to setup an easy approach to reproducibly induce commitment of healthy, transmission-competent gametocytes. We chose AlbuMAX as affordable and accessible reagent with reduced fatty acid content compared to human serum as potential inducer of sexual commitment.

We first tested if short exposure of ring stage parasites to media formulations containing 0.5% AlbuMAX induced gametocyte commitment and compared its performance with serum and recently reported minimal fatty acids (mFA) medium [16] as a high commitment induction control. Experiments were performed with cultured *P. falciparum* parasites (strain NF54 [17]) in an automated shaker system that changes the culture medium every 12 hours as previously described [18]. RPMI culture medium was supplemented with 10% human serum (Sanquin; Nijmegen, The Netherlands), 0.5% AlbuMAX (AlbuMAX II – Gibco 11021-045), or mFA media as described in [16]. Parasites were kept synchronized by enriching for late stage schizonts using Magnetic-Activated Cell Sorting (MACS) in the days prior to start of experiment. On day 0, cultures with parasites at 20-24 hours post infection (hpi) were seeded in 10ml shaker flasks at 1% parasitaemia and 5% erythrocytes (Sanquin; Nijmegen, The Netherlands). Parasites were exposed for 30 hours (from hereon referred to as ‘one day’) to AlbuMAX, mFA, or remained on normal serum media. In order to assess a single round of sexual commitment, at 30 hours post induction the media was changed in each condition to normal serum media with heparin (20U/mL) to block merozoite invasion of non-committed asexual parasites. This allows for commitment to be calculated by expressing the number of gametocytes formed as proportion of parasites that are present in the first cycle after induction. Blood smears were prepared and Giemsa-stained for parasitaemia (ring-stage parasites at 30 hours post induction) and gametocytaemia (gametocytes at day 5 post induction).

Gametocyte conversion rate in the automated shaker with the serum condition was 0.2% **(Figure 1A)**. For AlbuMAX supplemented media we observed an absolute increase in gametocyte conversion rate of 21.1% (95% CI: 8.1 – 34.0; p<0.001) and 50.1% (95% CI: 37.2 – 63.0; p<0.001) for mFA (**Figure 1A**). We repeated the assay in a 24 wells format using Nunc flat bottom plates as alternative culture system that is widely used, seeding 1ml culture per well with daily media change and plate incubation in a gassed humidified modular chamber (Billups and Rothenberg) at 37°C. Plates showed higher commitment rates overall with 6.5% commitment for the serum condition **(Figure 1B)**. However, there were similar trends when comparing the different induction conditions, with AlbuMAX and mFA supplemented media having higher (absolute difference) commitment compared to serum by 34.0% (95% CI: 21.5 – 46.5; p<0.001) and 63.2% (95% CI: 50.7 – 75.6; p<0.001), respectively **(Figure 1B)**. The higher commitment rate between both culture methods could be driven by a lower percentage of rings that were present post-induction **(Figure 1C and D)**; the number of rings was approximately halved when culturing in plates, reducing the denominator in the calculation of commitment rate, and could be due to the static growth conditions that are less optimal for parasite development compared to the shaker system. In addition, it is possible that the higher conversion rate for mFA media, i.e. the more depleted formulation, was a result of harsher growth conditions that resulted in a lower percentage of rings after induction compared to serum in both the automated shaker and plate **(Figure 1C and D)**.

**Figure 1:**
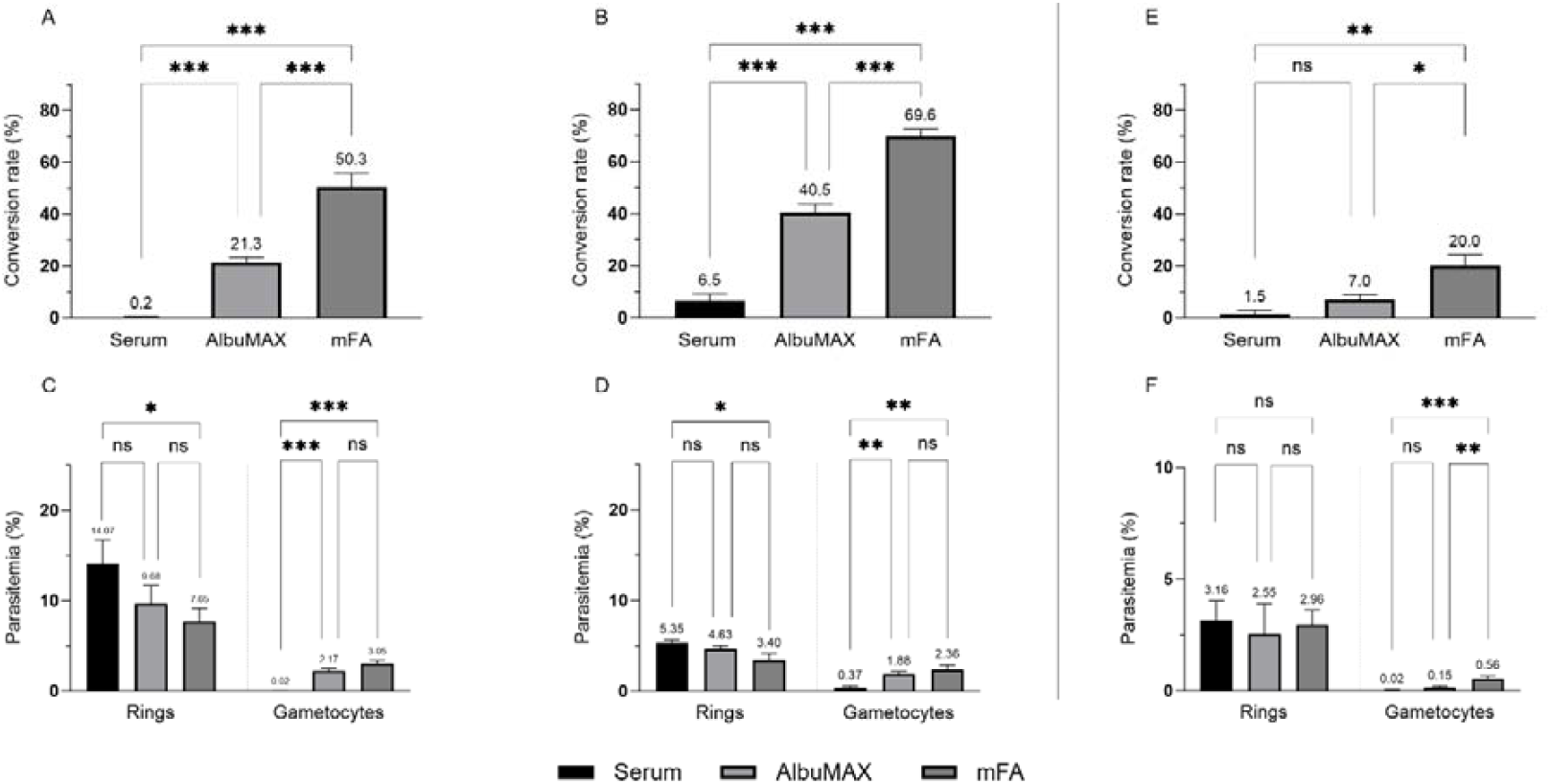
*P. falciparum* conversion from rings to gametocytes after exposure to media formulations. Bars show the conversion rate (%) with the standard error of the mean (SEM) for NF54 (**A-D**) or NF135 (**E-F**) ring stage parasites (20-24 hpi) that were cultured in the automated system (**A**) or in 24 wells plates (**B and E**). Serum supplemented medium was compared to single induction (1 day) of serum substitutes AlbuMAX or minimal fatty acids (mFA) to induce commitment. The conversion rate is the product (%) of committed rings to gametocytes, counted against 1000 erythrocytes (**C** for culture system, **D and F** for plates). Triplicate data from independent experiments are shown pooled. A linear regression was fit to compare the conversion rate between different conditions. No significance (ns): P > 0.05. Significance: * P ≤ 0.05, ** P ≤ 0.01 and *** P ≤ 0.001.

While the commitment rate for AlbuMAX gametocytaemia for both methods was similar. These results do show that AlbuMAX can be used for inducing *P. falciparum* sexual commitment. To test the possibility to induce gametocyte commitment in a genetically different parasite isolate (NF135, Asian origin [19]) we repeated the plate based conversion assay. While NF135 conversion rates were lower and more variable than NF54, we again observed a higher gametocyte conversion rate for AlbuMAX compared to serum, albeit not statistically significant (p=0.31) and highest conversion rates for mFA media (p=0.021 for comparison with serum; p=0.054 for comparison with AlbuMAX) **(Figure 1E and F)**, highlighting the potential broad use of the assay.

As the focus of this study was on AlbuMAX, which appears in these assays to be a more gentle formulation, and the total gametocyte load was not significantly different we continued to focus on exploring this media supplement for further gametocyte viability analysis post-induction in the commonly used transmission competent line NF54. For this we used exflagellation as a proxy for maturation and the transmission to mosquitoes as conclusive evidence of gametocyte quality. Transmissibility of gametocytes was assessed in membrane feeding assays using *An. stephensi* mosquitoes (Sind-Kasur strain) [18]. A standard approach to prepare for transmission assays is to use stress-induced conversion in serum supplemented media [18]; yielding workable but variable levels of commitment and transmission. To compare the AlbuMAX induction to this standard method, we used parasites cultured in the automated shaker system with serum media as a control. We tested one day exposure and also included four days AlbuMAX exposure to aim for increased gametocyte density by allowing a second and potential third cycle of asexual parasites to commit. No heparin was added in either condition to allow for ongoing replication and commitment. Parasites were evaluated between day ten and fourteen post induction for the ability of male gametocytes to activate (exflagellation) and transmission-competence to An. stephensi mosquitoes in the membrane feeding assay [18]. Exflagellation was induced with 50mM xanthurenic acid in serum supplemented media at room temperature with 15 minutes incubation before counting exflagellation centers on a haemocytometer; we observed an overall 14-fold (95% CI: 7.8 – 25.0; p<0.0001) increase for one day AlbuMAX and a 6.8-fold (95% CI: 3.7 – 12.0; p<0.0001) increase for four days AlbuMAX exposure compared to serum (**Figure 2A**). We further observed an overall increase of 33.5 fold (95% CI: 25.5 – 43.8; p<0.0001) and 2.8 fold (95% CI: 2.0 – 4.0; p<0.0001) in oocyst densities, respectively (**Figure 2B**). We did not observe evidence for an increase in gametocyte commitment in subsequent cycles by four days AlbuMAX exposure; instead, we observed 2.1-fold fewer exflagellation centers (95% CI: 1.2 – 3.7; p<0.0030) and 11.8-fold lower oocyst intensity (95% CI: 9.1 – 15.5; p<0.0001) for four versus one day of AlbuMAX. Superior performance of one-day AlbuMAX was particularly evident at days 10 and 11 post-induction (**Figure 2** and **Table 1**); prolonged AlbuMAX exposure may have resulted in reduced gametocyte viability related to sustained lower levels of fatty acids that are important for sexual parasites and play a critical role in formation of the membrane complex and gametocyte development [11]. Prolonged exposure to AlbuMAX may therefore be detrimental to early committed young stage gametocytes, even those within the first few days following commitment. This hypothesis appears supported by a delayed appearance of exflagellation centers and oocyst infection intensity (**Table 1**).

**Table 1:**
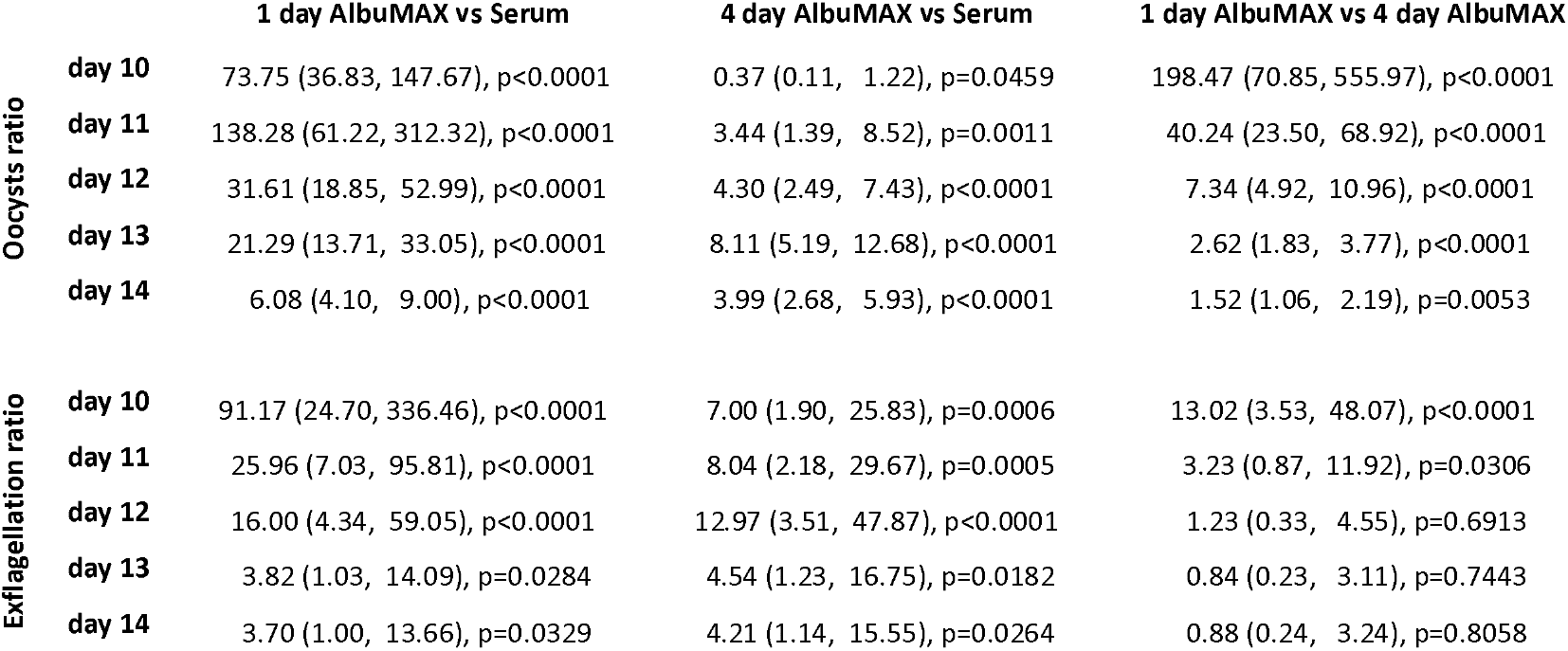
Ratio measures highlighting differences in conditions. Oocysts ratios were estimated using a negative binomial regression model and exflagellation ratios were estimated using a gamma regression model.

**Figure 2:**
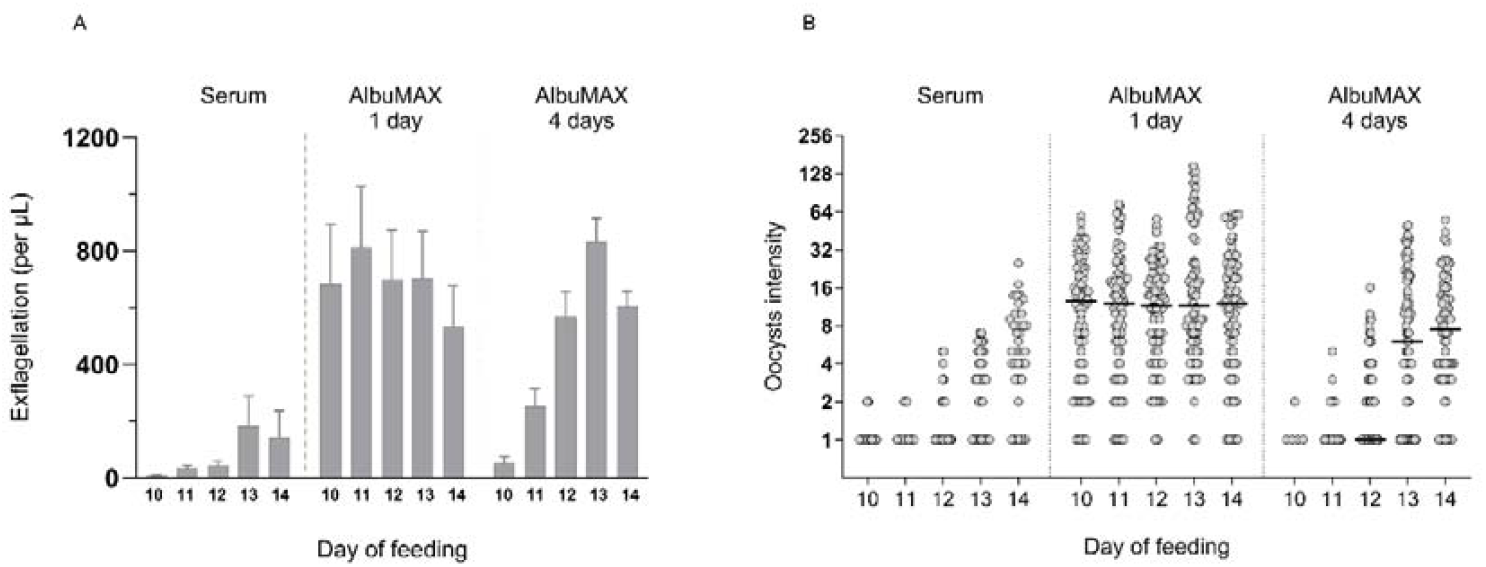
*P. falciparum* gametocyte commitment: serum compared to one or four days AlbuMAX induction. **A**. Bars show the number of exflagellation centers (per μL) with SEM evaluated on the days of mosquito membrane feeding. **B**. Scatter plot with dots representing oocyst intensity per mosquito for feeding between day 10 and 14 that were counted on day 7 post feeding, with bars displaying the median. A linear regression was fit to compare the conversion rate between different conditions. Triplicate data from independent experiments are shown pooled, statistical analysis is shown in **table 1**.

We conclude that one day exposure of *P. falciparum* cultured asexual stages to AlbuMAX supplemented media can consistently increase induction for commitment to gametocytogenesis with transmittable gametocytes being observed from day 10 onwards, but the prolonged exposure counteracts these gains. Clearly the timing of this approach is important, yet this also hints to the possibility of targeting fatty acids in early sexual stages as developmental controls. The presented induction methodology can easily be adopted by other culture facilities and allows a more stable and predictable production of gametocytes to be used in malaria (transmission) research.

## Declaration of Competing Interest

The authors report no declarations of interest.

### Project funding statement

This work was supported by an investment of Radboud University and Radboud university medical center. Teun Bousema and Wouter Graumans are further supported by a European Research Council (ERC) Consolidator Grant (ERC-CoG 864180; QUANTUM). The funders had no role in the design of experiments or the decision to publish findings.

## Acknowledgements

We would like to thank Jolanda Klaassen, Laura Pelser-Posthumus, Astrid Pouwelsen, Saskia Mulder and Jacqueline Kuhnen for their role in mosquito rearing and evaluation of membrane feeding experiments.

## Notes

### Competing Interest Statement

The authors have declared no competing interest.

### Summary of Updates

Figure 1 has been revised to include an additional P. falciparum isolate

